# EnrichMap: Spatially informed enrichment analysis for functional interpretation of spatial transcriptomics

**DOI:** 10.1101/2025.05.30.656960

**Authors:** Cenk Celik, Maria Secrier

**Affiliations:** UCL Genetics Institute, Department of Genetics, Evolution and Environment, University College London, Gower Street, London, WC1E 6BT, UK

## Abstract

Advances in spatial transcriptomics (ST) allow high-resolution mapping of gene expression within intact tissues, revealing spatially regulated biology. However, accurately quantifying gene set activity in spatial contexts remains inadequate. We present EnrichMap, a Python package designed to compute gene set enrichment scores in ST data by integrating batch correction, smoothing and spatial confounder adjustment. EnrichMap generates enrichment scores for signatures or pathways, with an optional framework for incorporating gene weights. To assess performance, we benchmark EnrichMap against established scoring techniques using spatial autocorrelation metrics such as Moran’s *I*. Our results show that EnrichMap improves spatial coherence, producing scores that capture both local and global spatial patterns, while conventional approaches tend to exhibit higher heterogeneity or diminished spatial structure. EnrichMap is demonstrated to be generalisable across a variety of ST platforms, including Visium, Xenium, MERFISH and imaging mass cytometry, maintaining consistent performance despite differences in spatial resolution and detection chemistry. Our method offers a platform-agnostic, spatially and biologically meaningful representation of gene set activity across complex tissue landscapes.

## BACKGROUND

The tissue microenvironment is a spatially organised ecosystem, comprising diverse cell types (e.g. epithelial, immune, stromal) and extracellular matrix components. These elements interact in structured spatial patterns shaped by development, homeostasis and pathology. In cancer, such interactions influence outcomes including immune evasion^1, 2^, metastasis^3, 4^ and therapeutic resistance^1, 5-7^. Advances in spatial transcriptomics (ST) enable transcriptome-wide profiling at near-cellular or cellular resolution, while preserving the spatial context^8, 9^. This spatially resolved view allows investigation of tissue *in situ*, where neighbouring cells share environmental cues and influence local phenotypes and cell states in cooperation. Capturing such intertwined spatial patterns is essential for understanding tissue function and diseases.

Gene set enrichment analysis (GSEA)^10^ and related approaches^11-14^ link expression data to biological processes. However, they assume independence between samples, an assumption violated in spatial data, where gene expression in a cell is often impacted by the surrounding microenvironment, accommodating the continuity of certain biological features across space. As a result, these methods fail to account for the spatial continuity of many biological features across tissue landscapes.

While several tools have been developed for domain detection (e.g. BayesSpace^15^, CellCharter^16^, SpottedPy^17^), spatial gene variability (e.g. SPARK^18^, SpatialDE^19^) or cell-cell communication (e.g. CellPhoneDB^20^, Renoir^21^, SpaceBF^22^), methods that integrate spatial context into the evaluation of gene programmes remain limited. Here, we present EnrichMap, a spatially informed framework for gene set scoring in ST data. EnrichMap integrates spatial relationships, models tissue continuity and corrects for spatial confounders by leveraging binary or weighted gene sets. The resulting maps reflect spatially organised signals, enabling discovery of biologically coherent tissue regions. EnrichMap also provides spatial diagnostics (e.g. autocorrelation, correlograms, variograms) adapted from geostatistics^23, 24^ to assess spatial structure.

We demonstrate EnrichMap’s utility across ST datasets, including healthy and cancer tissues. First, we showcase EnrichMap’s capabilities in a mouse brain dataset and recover a spatially localised pyramidal layer. Then, we investigate cancer hallmarks in two human breast cancer datasets to highlight EnrichMap’s batch handling. We demonstrate spatially localised proliferative zones and G0 arrest linked with immune suppression and metastasis, uncovering transcriptional landscapes that are not apparent from raw gene expression. Furthermore, we show EnrichMap’s performance across various ST platforms, including Visium HD, Xenium, MERFISH and imaging mass cytometry, highlighting its versatility in handling datasets with varying spatial resolution and detection chemistries. Our approach offers a platform-agnostic, spatially informed framework for functional genomics *in situ*, with wide-ranging applications in cancer research, developmental biology and tissue-based diagnostics.

## RESULTS

### EnrichMap enables spatially informed gene set enrichment analysis

Standard enrichment tools^10-14,25,26^, designed for either bulk or single-cell transcriptomics data, assume independent observations and overlook spatial relationships. When applied to ST data, these methods produce fragmented and artefactual patterns, limiting spatial interpretability. To address this, we developed EnrichMap, a spatially informed framework for gene set scoring in ST data. EnrichMap builds on the principle that biologically meaningful processes manifest as spatially coherent patterns across tissue sections by incorporating spatial smoothing, batch correction and spatial covariate adjustment (**Fig. 1a**), generating per-cell or per-spot enrichment scores visualised as interpretable maps with optional gene contribution plots (**Fig. 1b**).

**Fig. 1.**
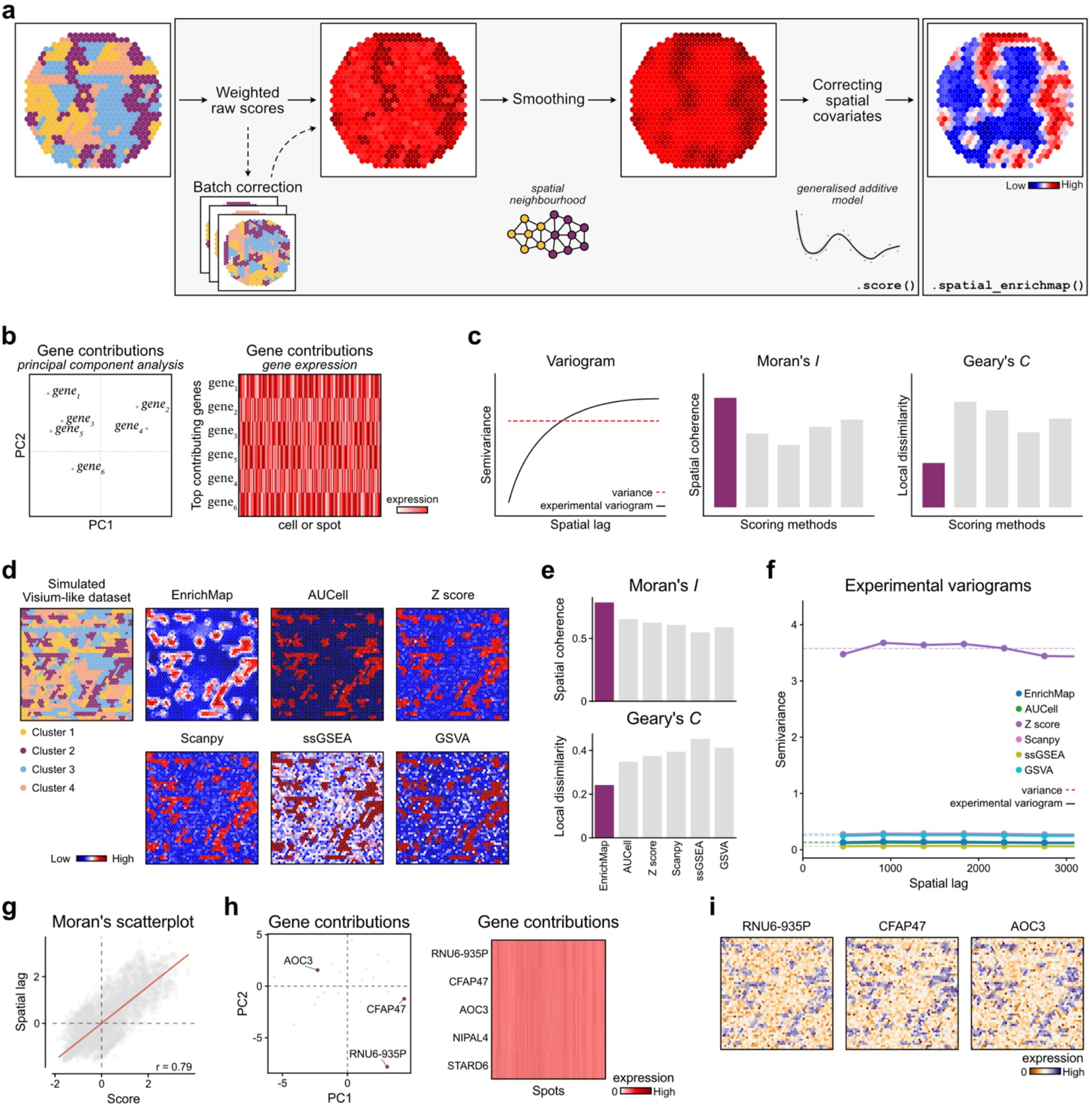
EnrichMap presents a spatially informed framework for gene set enrichment in spatial transcriptomics data. **a**, Overview of spatially aware gene set enrichment score workflow. EnrichMap calculates weighted raw scores, smoothens the scores based on spatial neighbourhood graph and corrects for spatial covariates, producing spatially informed enrichment scores. **b-c**, EnrichMap provides functionalities to identify the top contributing genes in a gene set, defined either by principal components (left) or by gene expression (right) **(b)**, and geostatistical analytical toolkit to aid the enrichment score’s spatial interpretability **(c). d**, EnrichMap showcased on a simulated Visium-like dataset, where a 10-gene signature was defined to identify cluster 2 and compared with conventional scoring methods, including AUCell, Z score, Scanpy, ssGSEA and GSVA. **e**, Spatial autocorrelation scores both in Moran’s *I* (top) and Geary’s *C* (bottom). **f**, Experimental variograms across different scoring methods (see Supplementary Fig. 1 for details). **g**, Moran’s scatter plot reveals a strong positive spatial autocorrelation, with a Moran’s *I* of 0.79, indicating that spatial units with high (or low) attribute values tend to be adjacent to others with similarly high (or low) values. This is evident from the dense clustering of points in the top-right (high-high) and bottom-left (low-low) quadrants, suggesting the scores exhibit significant spatial structuring. **h-i**, Top contributing genes **(h)** demonstrate consistent spatial coherence with their individual expression **(i)**.

The EnrichMap toolkit includes spatial diagnostics (i.e. Moran’s *I*, Geary’s *C*, variograms) to quantify spatial coherence and distinguish biological signal from technical noise (**Fig. 1c**), helping determine whether a gene programme shows meaningful spatial localisation. For instance, Moran’s *I* measures global spatial autocorrelation, capturing broad patterns where gene expression is spatially clustered, while Geary’s *C* is more sensitive to local dissimilarities, making it well-suited for detecting gradients or spatial boundaries. Similarly, a variogram shows how related (or dissimilar) two observations are as the spatial distance between them increases.

To demonstrate its utility, we applied EnrichMap to a simulated Visium-like dataset (**Fig. 1d**) with a 10-gene signature marking a specific population (cluster 2). Compared with area under the curve per cell (AUCell)^11^, Z score, Scanpy^27^ (`score_genes()`; Seurat’s `AddModuleScore()` function^26^), single-sample GSEA (ssGSEA)^12^ and gene set variation analysis (GSVA)^13^, EnrichMap yielded the most favourable values for both Moran’s *I* and Geary’s *C* (**Fig. 1e**), indicating superior sensitivity to spatial organisation. Unlike Z score, which produced spatially fragmented scores, EnrichMap preserved spatial coherence (**Fig. 1f, Supplementary Fig. 1a**). A Moran’s scatter plot confirmed strong spatial autocorrelation (Moran’s *I* = 0.79, *p*-value < 0.001; **Fig. 1g**), with high-to-high and low-to-low expression clustering across the tissue. Top contributing genes identified by EnrichMap showed spatial expression patterns that aligned with their enrichment scores (**Fig. 1h-i**), reinforcing the biological interpretability of the method. We further assessed sensitivity by scoring signatures of varying sizes. On the simulated data, signatures comprising 10-50 genes showed the most coherent spatial structure (**Supplementary Fig. 1b-d**), consistent with the gene set size of many co-regulated biological programmes. Repeating the simulation 100 times confirmed the robustness of these results, with EnrichMap consistently showing higher spatial autocorrelation compared with other methods, as captured by both variogram range distributions (**Supplementary Fig. 1e**) and Moran’s *I* scores (**Supplementary Fig. 1f)**.

### Spatially coherent gene programmes reveal laminar organisation in the mouse brain

To demonstrate EnrichMap’s capabilities in a real-world dataset, we explored a mouse coronal brain section dataset (**Fig. 2a**, Methods). The coronal section is highly compartmentalised, therefore, ideal to use for benchmarking. To test the performance of EnrichMap and other methods, we ranked the top 10 differentially expressed genes of the pyramidal layer, a layer of the cerebral cortex that contains large, excitatory pyramidal neurons responsible for integrating and transmitting cortical information.

**Fig. 2.**
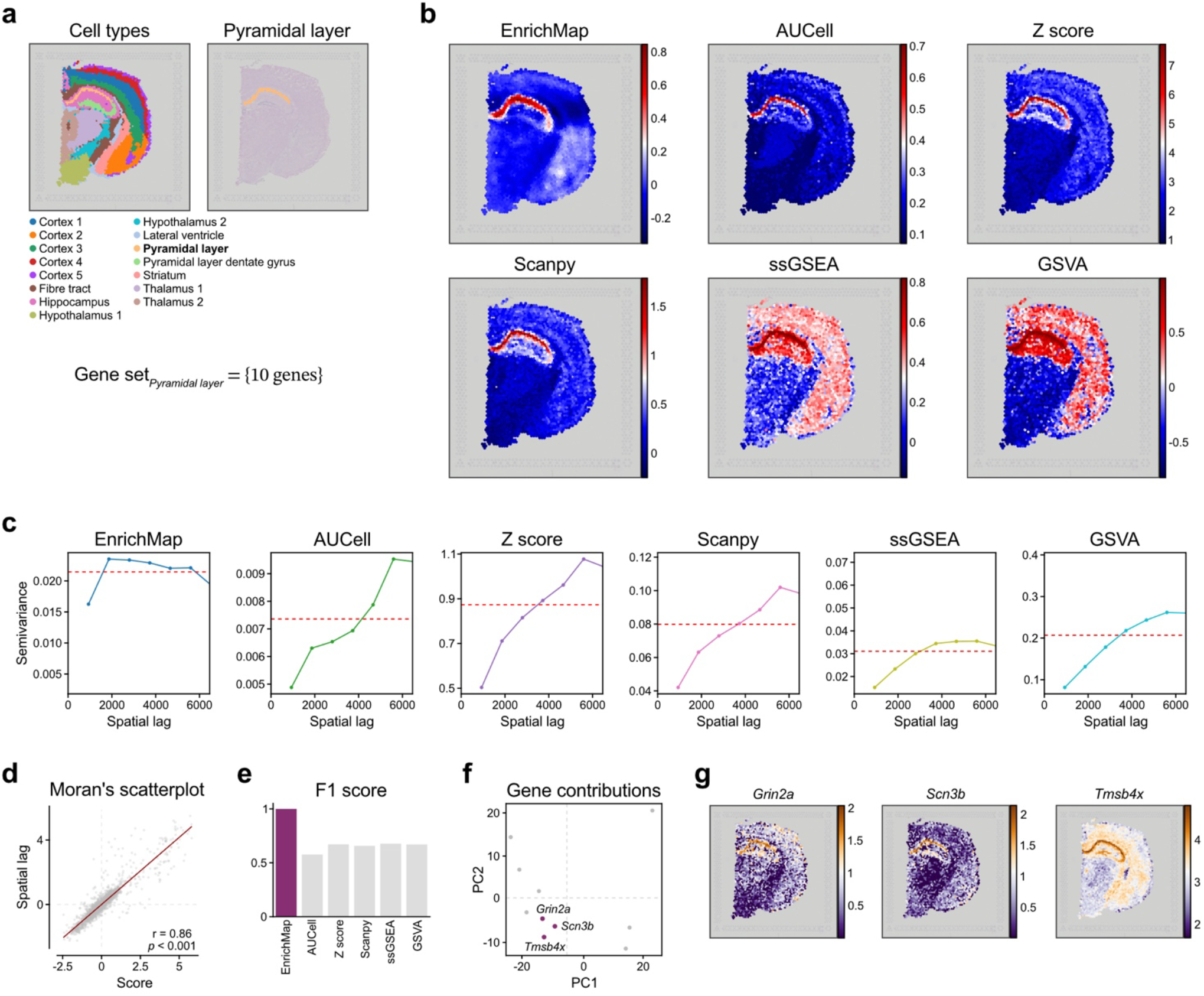
Laminar organisation in the mouse brain revealed by spatially informed gene set enrichment. **a**, An annotated Visium data set demonstrating structured anatomy of the mouse brain (left). A 10-gene signature was used to score for the pyramidal layer (right). **b**, EnrichMap scores demonstrate smoother tissue structure compared to AUCell, Z score, Scanpy, ssGSEA and GSVA. **c**, Variograms across different scoring methods highlight that EnrichMap has the best spatial structure, which increases sharply then plateaus, showing strong short-range structure, followed by ssGSEA. **d**, Moran’s scatter plot reveals a strong positive spatial autocorrelation of EnrichMap scores, with a Moran’s *I* of 0.86 with *p* < 0.001. **e**, Bar plot shows F1 scores across different scoring methods in identifying the pyramidal cells. EnrichMap outperforms all other methods, achieving the highest score. **f-g**, EnrichMap finds the genes *Grin2a, Scn3b* and *Tmsb4x* as top contributors that clustered together **(f)**, aligning with their individual expression **(g)**.

EnrichMap demonstrated the smoothest spatial structure of the pyramidal layer scores both visually (**Fig. 2b**) and statistically, where a sharp increase was followed by a plateau in the variogram analysis (**Fig. 2c**). To enable a fair comparison across methods, we did not centre the generated scores, although EnrichMap inherently produces scores centred around zero, reflecting both activation and deactivation of the input gene set. Nonetheless, Z score and AUCell were spatially unstructured as observed in the constant increase of the corresponding variograms, whereas Scanpy, ssGSEA and GSVA remained spatially autocorrelated to some extent (**Supplementary Fig. 2a**) and EnrichMap yielding autocorrelation of a Moran’s *I* of 0.86 with a *P* value of 0.001 (**Fig 2d**).

To benchmark classification accuracy, we evaluated each method’s ability to recover a predefined cell population. EnrichMap achieved the highest classification performance (F1=1.0), significantly outperforming all other methods (**Fig. 2e**) that showed comparable but consistently lower performance (F1 ≈ 0.6-0.7). These results underscore EnrichMap’s sensitivity and precision in capturing spatially meaningful biological programmes, supporting for spatially resolved functional inference. We further compared runtimes across methods using gene sets of varying size. Z score and Scanpy were the fastest, followed by AUCell and ssGSEA, with GSVA being the slowest (**Supplementary Fig. 2b**). EnrichMap showed stable and relatively short runtimes up to 100 genes, after which a linear increase was observed, reflecting the cost of neighbourhood-based calculations–a feature that conventional methods ignore. Nevertheless, all methods completed within seconds, rendering runtime differences negligible in practice.

We next explored the top contributing genes and found that *Grin2a, Scn3a* and *Tmsb4x* were among the highest contributors that clustered together (**Supplementary Fig. 2c**), highlighting the deep layer, involved in modulating excitatory circuits in the brain, the main roles of pyramidal neurons in CA1 region of the hippocampus^28^.

### Design considerations for accurate spatial enrichment with EnrichMap

EnrichMap implements a coefficient of variation (CV)^29^-based gene weighting scheme which prioritises genes with high relative dispersion to enhance sensitivity to biologically relevant, spatially structured signals (**Supplementary Fig. 3a-b**). To explore this, we assessed the influence of housekeeping genes, whose expression is typically ubiquitous across tissues regions. When housekeeping genes were added to the pyramidal layer signature, EnrichMap continued to capture spatially coherent structures, yielding Moran’s *I* scores comparable to those without housekeeping genes and showing high rank correlation with the original scores (**Supplementary Fig. 3c-d**). This demonstrates that EnrichMap scores remain robust and spatially coherent, even in the presence of uniformly expressed genes, thereby mitigating noise from genes with weak or no spatial structure.

We evaluated EnrichMap’s sensitivity to a range of gene set sizes. Very small signatures (e.g. 2 or 5 genes) produced patchy enrichment patterns, whereas larger signatures (200-500 genes) led to overestimation (**Fig. 3a**). These findings suggest that signatures comprising 10-50 genes offer not only spatially but also biologically relevant balance. While variogram shapes were consistent across all sizes, semivariance was most stable within the 10-50 gene range (**Fig. 3b**). Notably, Moran’s *I* was highest at 50 genes, indicating the strongest spatial autocorrelation (**Fig. 3c, Supplementary Fig. 4a-b**). Although peak autocorrelation was observed with 50 genes, gene sets of 10-20 or even 100 genes may still be appropriate, as their enrichment scores correlate with those from 50-gene signatures at over 93% accuracy (**Supplementary Fig. 4c**). From here onwards, we centred all spatial plots according to EnrichMap’s design to ensure consistent and interpretable visualisation. We also provide a wrapper function for generating spatial enrichment maps in an interpretable format.

**Fig. 3.**
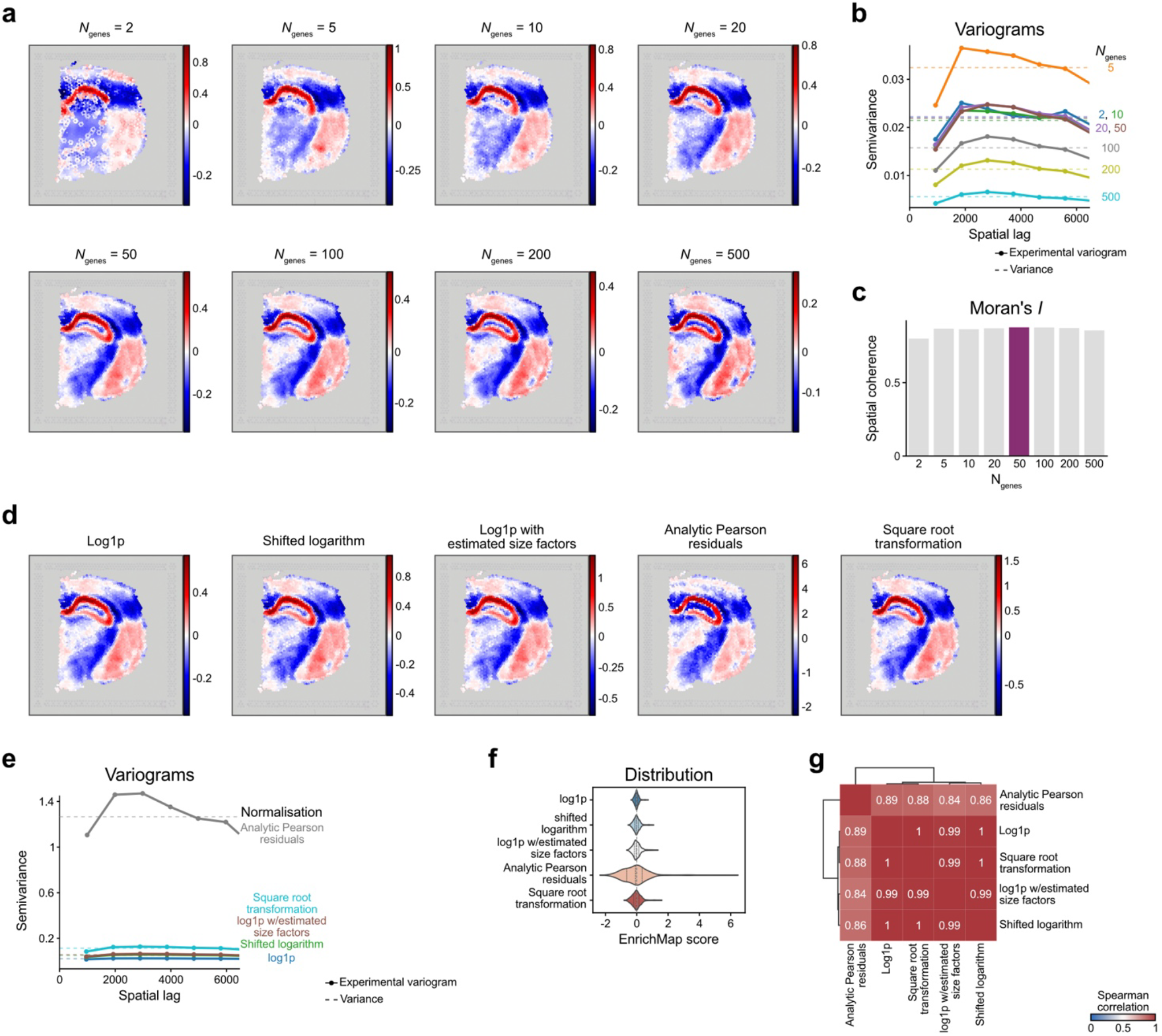
EnrichMap produces consistent enrichment scores across gene set sizes and normalisation methods. **a**, Spatial enrichment scores generated by EnrichMap using gene sets of varying sizes. **b**, Variograms across gene set sizes demonstrate stability for signatures comprising 10–50 genes. **c**, Moran’s *I* indicates peak spatial autocorrelation with a 50-gene signature. **d**, Enrichment scores derived using different normalisation methods. **e**, Variograms remain stable across normalisations, except for analytic Pearson residuals, which show greater variability. **f**, Distribution of enrichment scores across normalisation methods. **g**, Similarity matrix showing high concordance (>99%) among most normalisation methods.

We then explored the effect of neighbourhood size on spatial associations. Using 1-2 neighbourhood rings (i.e., 6-12 neighbours) most accurately highlighted the ground truth pyramidal layer without inflating enrichment scores. Both neighbourhood ring analysis and variogram-based assessment indicated that neighbourhood size should be predefined to avoid artificial score inflation (**Supplementary Fig. 3e-f**). This adaptability makes EnrichMap well suited to platforms like Visium, where spatial resolution and spot topology can vary.

Although EnrichMap can operate directly on raw counts (**Supplementary Fig. 4d**), we recommend applying normalisation to improve smoothness and mitigate technical artefacts, such as those introduced by batch effects. To evaluate this, we compared EnrichMap’s performance using several commonly used approaches: shifted logarithm transformation^30^, natural logarithm of one plus (log1p) with estimated size factors^31^, analytic Pearson residuals^32^, square root transformation and standard log1p normalisation (see Methods). Apart from analytic Pearson residuals, all methods yielded consistent results (**Fig. 3d-f**), with spatial enrichment scores exhibiting over 99% similarity (**Fig. 3g**). Analytic Pearson residuals, although still producing interpretable results, showed greater variability in spatial score distributions (**Fig. 3e-f, Supplementary Fig. 4e**). These findings support the use of log-based transformations for robust scoring.

### EnrichMap reveals spatially distinct cancer hallmark programmes in breast tumours

To demonstrate EnrichMap’s broader applicability beyond developmental contexts, we investigated the spatial organisation of cancer hallmarks in breast cancer. Using two ST slides from distinct patients, we applied previously established signatures^33, 34^ (see Methods) to explore proliferation and cell cycle arrest patterns within these cancer tissues using EnrichMap (**Fig. 4a**). Proliferation and G0 arrest scores were computed within tumour-annotated spots only (**Fig. 4a**). Despite the spatial the restriction to tumour-only regions, the resulting enrichment maps exhibited clear spatial patterns, smooth transitions and high spatial autocorrelation in both slides (**Fig. 4b-c**). As expected, G0 arrest and proliferation scores were inversely correlated, reflecting their spatially mutually exclusive biological states (**Fig. 4b**), reinforcing EnrichMap’s reliability in recapitulating spatial cell cycle dynamics.

**Fig. 4.**
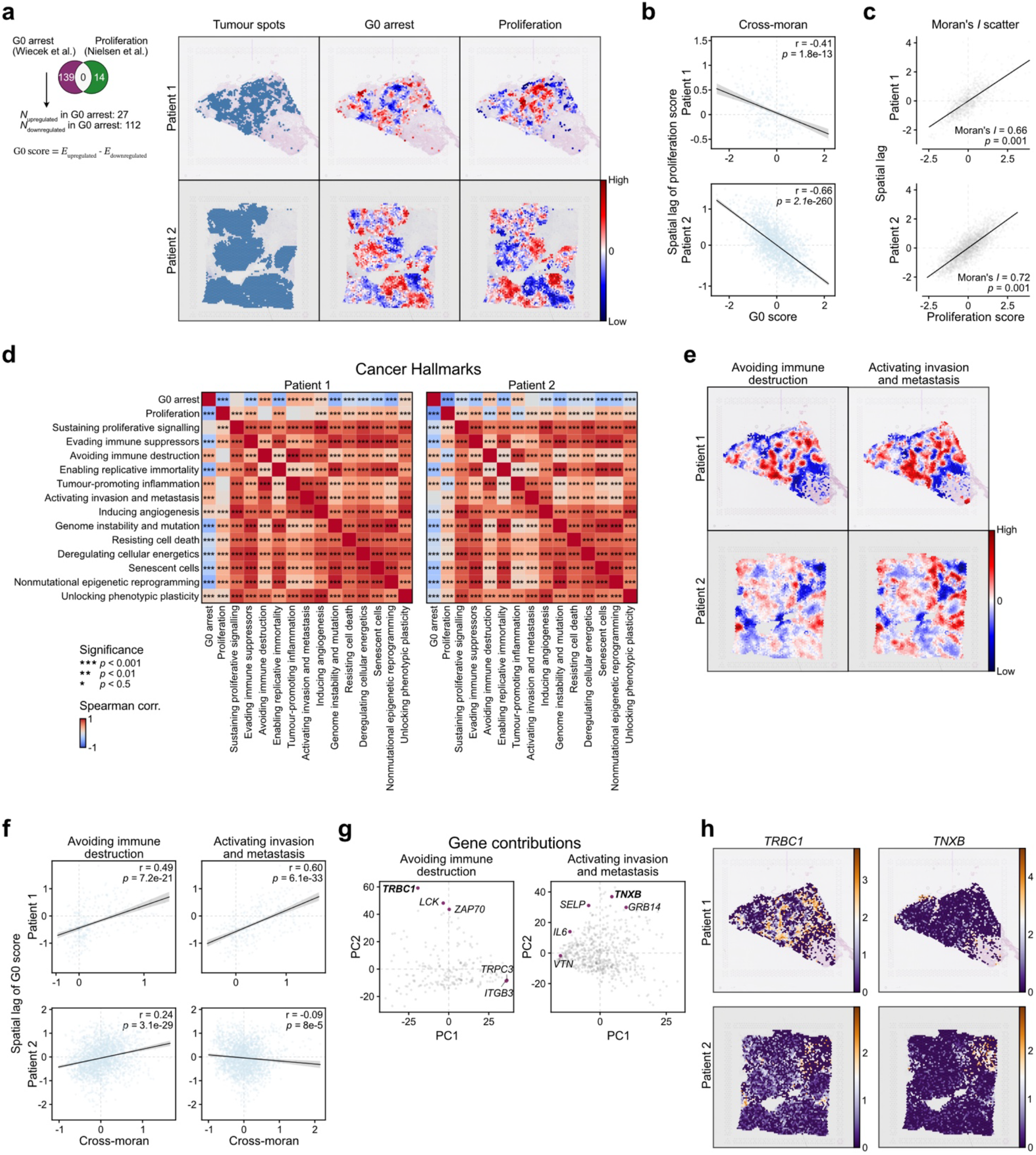
EnrichMap reveals spatially distinct G0 arrest niches linked to immune evasion and metastasis. **a**, Spatial G0 arrest (Wiecek *et al*.) and proliferation scores (Nielsen *et al*.) computed using EnrichMap in two breast cancer patients, restricted to tumour-annotated spots. **b**, Cross-Moran of proliferation enrichment scores reveal expected spatial anticorrelation, indicating mutually exclusive localisation of quiescent and proliferative tumour niches. **c**, Moran’s *I* of proliferation scores demonstrate consistent and smooth spatial patterns in both patients. **d**, Cancer hallmark signatures avoiding immune destruction and tumour-promoting inflammation show correlation with G0 arrest across both patients. **e-f**, Spatial map of EnrichMap scores **(e)** and cross-Moran correlation with *avoiding immune destruction* and *activating invasion and metastasis* hallmarks **(f)** across two patients. **g-h**, Gene-level analysis of hallmark signatures contributing to G0-enriched niches show top 5 genes **(g)** and gene expression of the selected genes **(h)**.

We next assessed the spatial association of G0 arrest and proliferation with other cancer hallmarks using curated signatures from Sibai et al. (2025)^35^. Proliferation co-localised with hallmarks such as *sustaining proliferative signalling, resisting cell death* and *replicative immortality* (**Fig. 4d-e, Supplementary Fig. 5a**). In contrast, G0 arrest was consistently co-enriched with *avoiding immune destruction* and *tumour promoting inflammation* hallmarks, suggesting G0 regions may represent immune-suppressed niches whose features have been linked with metastatic progression^36, 37^ and therapy resistance^38-40^. Notably, in patient 1, G0 arrest also aligned with *activation of invasion and metastasis*, a pattern absent in patient 2 (**Fig. 4d-f**), underscoring interpatient heterogeneity and EnrichMap’s capacity to capture heterogeneous spatial programmes across individuals. To explore the molecular drivers these programmes, we examined gene-level contributions to immune evasion and metastasis (**Fig. 4g–h, Supplementary Fig. 5b–c**). While overarching spatial patterns were conserved, the specific gene-level contributors varied between patients, revealing interpatient variability in the expression of genes *TRBC1* and *TNXB* (**Fig. 4h**). Together, these findings highlight EnrichMap’s ability to detect spatially distinct, biologically meaningful programmes, with potential implications for diagnostics and spatially targeted therapies.

### Platform-agnostic spatial enrichment with EnrichMap

To assess the generalisability of EnrichMap across diverse spatial transcriptomics (ST) platforms, we applied it to datasets spanning a range of spatial resolutions, detection chemistries and gene capture strategies. These included high-resolution Visium HD (10x Genomics), Xenium (10x Genomics), MERFISH (Vizgen) and imaging mass cytometry (Standard Bio), each offering distinct advantages in spatial granularity, throughput and tissue compatibility.

In a coronal mouse brain section profiled using Visium HD, which provides denser spatial coverage than standard Visium, EnrichMap generated smooth enrichment landscapes using the same gene signature applied to standard Visium data (**Fig. 5a**). The top contributing genes exhibited spatial patterns that cohered with known anatomical structures and closely mirrored the EnrichMap scores.

**Fig. 5.**
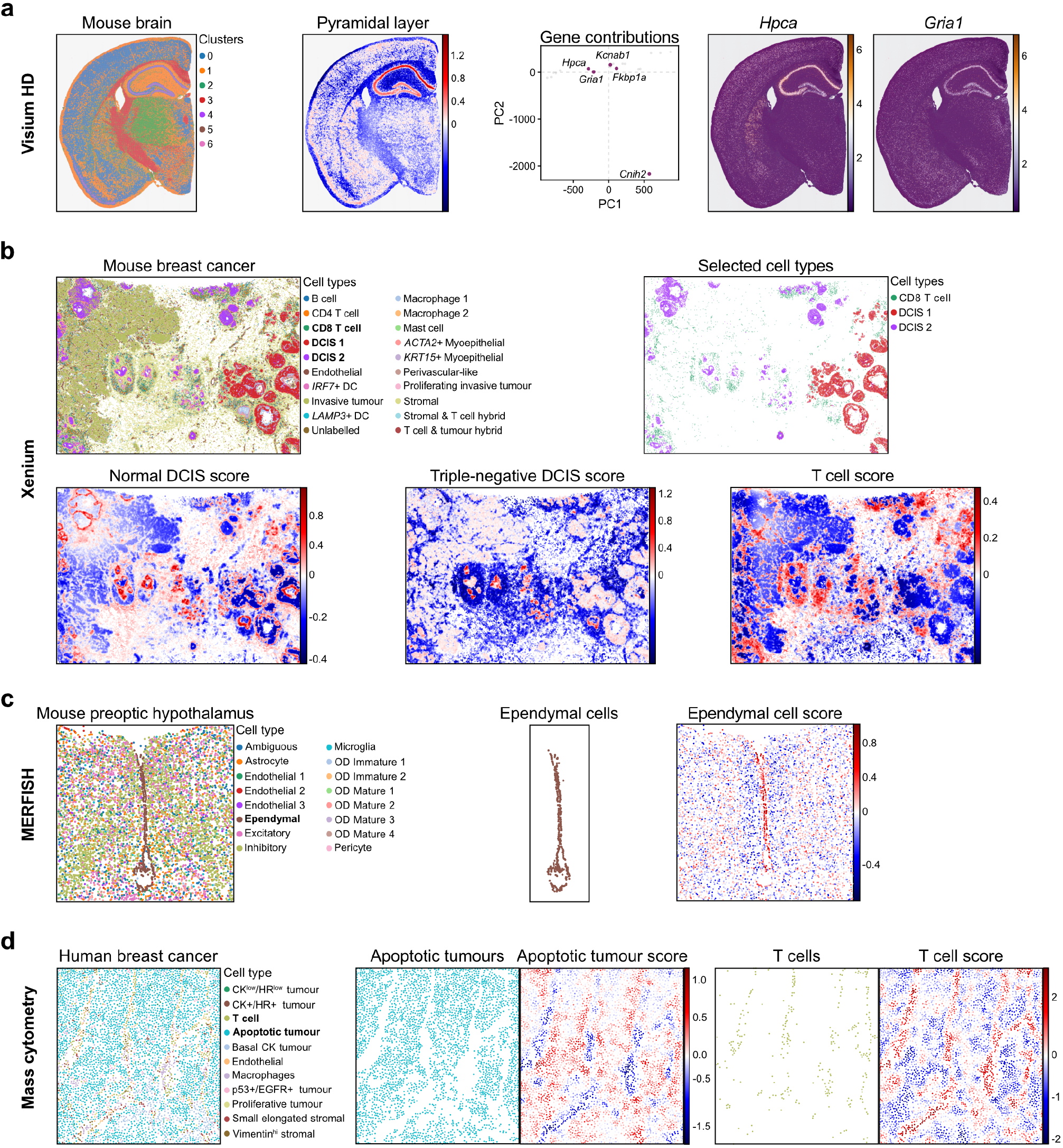
EnrichMap generalises across spatial transcriptomics platforms and resolutions. **a**, Application of EnrichMap to the Visium HD mouse brain dataset. Left: spatial clusters; middle: EnrichMap score for the pyramidal layer; right: top gene contributors with expression patterns for two selected genes. **b**, Annotated mouse breast cancer Xenium dataset (top left) and targeted cell types (top right). Bottom panels show EnrichMap scores for Normal DCIS (bottom left), triple-negative DCIS (bottom middle), and T cells (bottom right). **c**, Mouse preoptic hypothalamus MERFISH dataset with tissue annotations (left), ependymal cell regions highlighted for clarity (middle), and EnrichMap score for ependymal cells (right). **d**, Mass cytometry of human breast cancer. Left: spatial context; middle: apoptotic tumour regions captured with two antibodies; right: T cells identified using three antibodies.

In a Xenium dataset of mouse breast cancer tissue, EnrichMap retained its capacity to resolve fine-grained enrichment patterns at single-cell resolution. It identified spatially distinct niches corresponding to normal ductal carcinoma *in situ* (DCIS), triple-negative DCIS and T cells (**Fig. 5b**), despite the increased sparsity and noise characteristic of single-cell resolved spatial data.

In the MERFISH mouse preoptic hypothalamus dataset, which captures high-resolution expression for a targeted gene panel, EnrichMap successfully identified enrichment of ependymal cell signatures in line with original tissue annotations (**Fig. 5c**). This result underscores EnrichMap’s robustness in settings with constrained gene coverage.

Finally, to evaluate its applicability to spatial proteomics, we applied EnrichMap to an imaging mass cytometry dataset^41^. Despite the minimal input of only two or three antibodies, EnrichMap captured distinct enrichment patterns associated with apoptotic tumour cells and T cells (Fig. 5d), suggesting its potential utility in low-dimensional spatial data modalities. However, this approach is best suited to cases where each cell type or biological process is uniquely identifiable by a small, non-overlapping set of markers, as overlapping expression patterns could limit the interpretability of signals in sparse antibody panels.

Together, these results demonstrate that EnrichMap is platform-agnostic and broadly applicable across diverse spatial omics technologies. Its ability to extract coherent biological signals in both transcriptomic and proteomic data, regardless of resolution or gene coverage, highlights its utility for integrative spatial analysis across experimental platforms.

## DISCUSSION

The spatial organisation of biological function defines tissue biology, and in diseases like cancer, underpins tumour evolution, immune evasion and treatment response, yet the analytical tools to extract spatially coherent insights remain limited. Here, we introduced EnrichMap, a method that bridges this gap by enabling spatially aware GSEA directly on ST data.

EnrichMap addresses a fundamental shortcoming of conventional GSEA methods: the assumption of sample independence. By explicitly modelling spatial context and continuity, EnrichMap produces scores that reflect not only gene expression magnitude but also the spatial structure of underlying biological processes. Through spatial smoothing and spatial confounder correction, the method yields functional maps that better correspond to known histological compartments and tissue architecture. Our analysis of human breast cancer tissues reveals patterns of immune suppression, proliferative gradients and invasive activity that would be missed using classical enrichment approaches. For example, Sibai et al. (2025)^35^ observed that proliferative signalling co-localised spatially with hallmarks such as *evading growth suppressors, senescent cells* and *non-mutational epigenetic reprogramming* across multiple cancer types, patterns that were also captured by EnrichMap. Notably, while their model failed to detect coherent regions associated with *avoiding immune destruction* and *tumour-promoting inflammation*, EnrichMap successfully identified these signals, particularly within G0 arrested regions. These findings highlight EnrichMap’s capacity to detect spatially coherent biological signals beyond the reach of conventional methods, offering a powerful framework for spatially informed functional genomics.

A distinguishing feature of EnrichMap is its inclusion of spatial diagnostics, which allow users to assess whether inferred gene set activity exhibits expected spatial continuity. These tools not only strengthen confidence in results but also provide a means to detect potential artefacts or spatial biases in the data. EnrichMap thereby encourages a more principled and statistically grounded interpretation of spatial gene programmes. For instance, from all our analyses, we found that using 10-50 genes, and even up to 100 genes, demonstrates spatial coherence.

Compared to other ST tools, which primarily focus on spatial domain detection, spatially variable genes or cell-cell communication, EnrichMap fills an important niche by offering spatial gene programme scoring. While tools like SpatialDE^19^ and SPARK^18^ identify where genes vary or cluster, they do not assess coordinated expression of gene sets within a spatial framework. By doing so, EnrichMap supports a richer layer of biological interpretation that is especially valuable in complex diseases such as cancer, where distinct phenotypes arise through the interplay of multiple cell types and signalling pathways within spatially organised niches.

Nonetheless, EnrichMap has several limitations. The current framework is designed for 2D ST data and may need adaptation for 3D or multi-slice technologies. Second, spatial smoothing improves noise robustness but can blur sharp boundaries in low-resolution data, making it imperative to assess spatial metrics such as variograms and autocorrelation. While EnrichMap supports user-defined gene weights, finding optimal weights remains a challenge. We addressed this by implementing the coefficient of variation (CV)^29^ as a simple heuristic to prioritise genes with high relative dispersion, which often reflect spatially patterned, biologically relevant expression. Low-CV genes, such as housekeeping genes, contribute less to spatial variation. Applying EnrichMap to a 10-gene pyramidal layer signature, a housekeeping gene set, and a combined signature showed CV-based weighting correctly prioritised spatially specific genes, though some housekeeping genes (e.g. *B2m*) still ranked highly but lacked spatial specificity. While more complex weighting strategies exist, CV offers an efficient, unsupervised proxy without requiring prior knowledge. To demonstrate this, we applied EnrichMap to the 10-gene pyramidal layer signature (Fig. 2a), a housekeeping gene set^42^ and a combined signature (Supplementary Fig. 3a-d), and CV-based weighting correctly prioritised the pyramidal layer genes, even in the mixed set. Notably, *B2m*, a housekeeping gene, ranked the last among the top contributors but clustered with other housekeeping genes, indicating low spatial specificity, and stability as reported by Ho and Patrizi (2021)^42^. These results demonstrate both the usefulness and limitations of CV, highlighting the need for contextual interpretation of gene weights.

We also envision several exciting directions. Integration with cell-cell interaction frameworks could allow gene set scoring to be modulated not only by spatial proximity but also by inferred communication networks. EnrichMap could also serve as a prior for spatial deconvolution methods, helping refine cell type annotations based on pathway activity. Moreover, extending the framework to incorporate temporal or multi-modal data, such as spatial proteomics or metabolomics, could provide an even more comprehensive view of tissue organisation.

In conclusion, EnrichMap provides a spatially aware, interpretable and extensible framework for functional analysis of ST data. By capturing the continuity and structure of gene programme activity across tissues, it offers a powerful approach for spatial functional genomics, particularly in the context of cancer, where tissue architecture and microenvironmental context are critical determinants of disease behaviour and therapeutic response.

## METHODS

### EnrichMap: Spatially aware gene set enrichment scoring

To compute gene set enrichment scores that incorporate spatial context and account for batch effects, we developed EnrichMap, a Python tool compatible with scverse ecosystem, including `anndata, scanpy` and `squidpy`. EnrichMap supports multiple gene set signatures, offers optional spatial smoothing and corrects for spatial covariates using generalised additive models (GAM), accommodating the continuity of certain biological features across space.

#### Raw gene set score computation

Let *x*_*ij*_ denote the expression of gene *g*_*i*_, and *w*_*i*_ the weight assigned to a gene *g*_*i*_. The raw score *Z*_*j*_ for each cell or spot is computed as a weighted average over the genes:

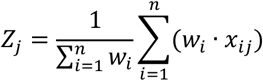

A dictionary of gene weights may be provided for the gene signature *S* = {*g*_*1*_, *g*_2_, …, *g*_3_} or alternatively inferred from the expression matrix (see next section). By default, EnrichMap infers gene weights from the normalised expression values of the signature genes in the matrix (i.e. `adata.X`). When a batch label (i.e. representing multiple slides) is specified via `batch_key`, z-score normalisation is applied within each batch. For a given batch *b*, batch-normalised scores are computed as:

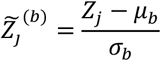

where *µ*_*b*_ and *σ*_*b*_ are the mean and standard deviation of raw scores within batch *b*, respectively.

#### Spatial smoothing

To incorporate local spatial context, we construct a `spatial neighbours’ graph leveraging squidpy’s spatial_neighbors()` function. For each cell or spot, the smoothed score 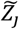, is calculated as a weighted average of the scores of its neighbours:

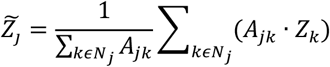

where *A* is the adjacency matrix of the spatial graph and *N*_(*j*)_ denotes the neighbours of the cell *j*.

#### Correcting spatial covariates

To account for spatial trends in gene expression, we fit a generalised additive model (GAM) using spatial coordinates as smooth covariates:

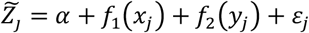

Here, *f*_1_ and *f*_2_ are spline functions of the *x* and *y* spatial coordinates, respectively, *α* is the intercept and ε_*j*_ is the residuals. The corrected score is then taken as the model residual:

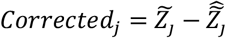

All steps are integrated into the `score()` function available under `enrichmap.tools` module. The final corrected scores for each signature are stored in `adata.obs`. We also implemented a wrapper function `spatial_enrichmap()` for intuitive visualisation of scores under `enrichmap.plotting` module.

### Inferring gene weights by the coefficient of variation

To infer gene weights, EnrichMap uses the coefficient of variation (CV)^29^. For each gene *g*_*i*_, we compute the mean *µ*_*i*_ and standard deviation *σ*_*i*_ of its expression across all cells or spots. The coefficient of variation is defined as:

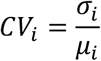

For genes where *µ*_*i*_ = 0, the CV is set to zero to avoid division by zero. The result is a dictionary *W*mapping gene names to their inferred weights:

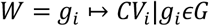

This scheme captures relative variability and upweights genes with higher dynamic expression, which may highlight spatially contextual gene expression patterns more robustly than uniform weighting.

### Other functionalities

For interpretation of the spatial gene set enrichment scores, we developed several visual and analytical tools. For example, the `gene_contributions_heatmap()` function visualises the expression of the top contributing genes to any given enrichment score across all data points in a slide. Genes are sorted by expression level, and the heatmap highlights which genes drive specific spatial patterns across conditions or samples. Similarly, to explore broader patterns in gene contributions, we implemented the `gene_contributions_pca()` function, which applies principal component analysis (PCA) to reduce the dimensionality of gene-level weights of a given enrichment score. This approach facilitates the identification of dominant contribution patterns and gene clusters.

Spatial autocorrelation of enrichment scores was assessed using the `morans_correlogram()` function. This method computes Moran’s *I* statistic over increasing neighbourhood distances, generating a spatial correlogram that captures how spatial structure changes with distance. The implementation draws on geospatial statistics functions from the `esda` and `libpysal` libraries^23,24^.

Finally, spatial continuity and heterogeneity can be evaluated using the `variogram()` function, which plots a semi-variogram for a given enrichment score. This classical geostatistical tool estimates how variance evolves with increasing spatial lag and relies on functionality from the `esda` and `libpysal` libraries^23, 24^.

### Spatial transcriptomics datasets

All tests of the EnrichMap package were performed using a pre-processed and annotated coronal section of the mouse brain dataset provided by the `squidpy` package^43^. The raw version of this dataset is publicly available at https://support.10xgenomics.com/spatial-gene-expression/datasets. To demonstrate cross platform compatibility of EnrichMap, we additionally tested EnrichMap on three further datasets: a MERFISH dataset from Moffitt et al (2018)^44^, and Visium HD mouse brain and Xenium mouse breast cancer datasets, also available from https://support.10xgenomics.com/spatial-gene-expression/datasets.

As the datasets were provided with raw counts, we first filtered for genes with at least 1 count in a single spot. Following this, we applied `scanpy’s` normalisation and log1p transformation, except in analyses specifically designed to evaluate the impact of different normalisation strategies on EnrichMap scores. In those cases, we compared standard log(*X* + 1) (log1p) transformation with alternative approaches including pooling-based size factor estimation (log1p with estimated size factors)^31^, shifted logarithm 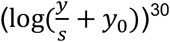, analytic Pearson residuals 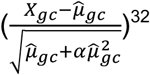 and square root transformation 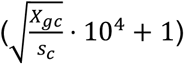.

### Simulated spatial transcriptomic dataset

A hexagonal grid mimicking 10X Visium layout was generated with fixed spacing to simulate spatial coordinates. Human gene symbols (n = 15,000) were sampled from the HGNC database [https://www.genenames.org]. Spatially coherent clusters were assigned by randomly sampling biological labels and reinforcing local neighbourhood similarity. Gene expression was simulated using a Poisson distribution, with elevated expression for a random gene set in cluster 2. The resulting data were stored in an AnnData object with spatial coordinates.

### Gene set signatures

For the Visium mouse brain dataset, differentially expressed genes (DEGs) were identified from selected clusters using a minimum log_2_ fold change of 1 and a detection fraction of 0.5. The top 10 genes were then used for scoring on the Visium and 50 genes for the Visium HD mouse brain dataset. The same strategy was applied to the simulated dataset. To assess spatial metrics and performance, we generated gene sets of varying sizes by selecting the top 2, 5, 10, 20, 50, 100, 200 and 500 genes from the DEG lists. Additionally, a list of mouse housekeeping genes was used to illustrate the impact of gene weighting^42^.

For the Xenium mouse breast cancer data set, we employed two signatures for identifying triple negative ductal carcinoma *in situ* (DCIS) and normal DCIS regions reported by Rebbeck et al. (2022)^45^. We also used an 8-gene cytotoxic T cell signature^46^ to highlight T cells across invasive and non-invasive regions of the breast tumour.

The G0 arrest signature was calculated by EnrichMap generated scores of 27 upregulated and 112 downregulated genes described by us in Wiecek et al (2023)^47^. A final G0 arrest score was obtained by subtracting the two scores. A unique proliferation signature^34^ was then used for validation. Cancer hallmark gene sets were generated as described in Sibai et al (2025)^35^.

## Data availability

The mouse brain coronal section (Visium), mouse preoptic hypothalamus (MERFISH) and human breast cancer (imaging mass cytometry) datasets are available through the `squidpy` package^43^. Mouse breast cancer (Xenium) data can be accessed via 10x Genomics at https://support.10xgenomics.com/spatial-gene-expression/datasets. Processed Visium from human breast cancer samples and Visium HD mouse coronal brain section data are publicly available on Zenodo (DOI: 10.5281/zenodo.15438169).

## Code availability

EnrichMap is a pip installable Python package available at https://github.com/secrierlab/enrichmap, with documentation at https://enrichmap.readthedocs.io/en/stable. All the code to reproduce the results of the analyses can be found at the following repository: https://github.com/secrierlab/enrichmap_reproducibility.

## Acknowledgements

MS and CC were supported by a UKRI Future Leaders Fellowship (MR/T042184/1, MR/Y034031/1). Work in MS’s lab was supported by a BBSRC equipment grant (BB/R01356X/1) and a Wellcome Institutional Strategic Support Fund (204841/Z/16/Z). We thank Ms Emma Champneys for her critical reading of the manuscript.

## Author’s contributions

CC conceptualised and conducted the study. MS supervised the study and secured the funding. CC wrote the manuscript with input from MS.

## Supplementary Figures

**Supplementary Fig. 1.**
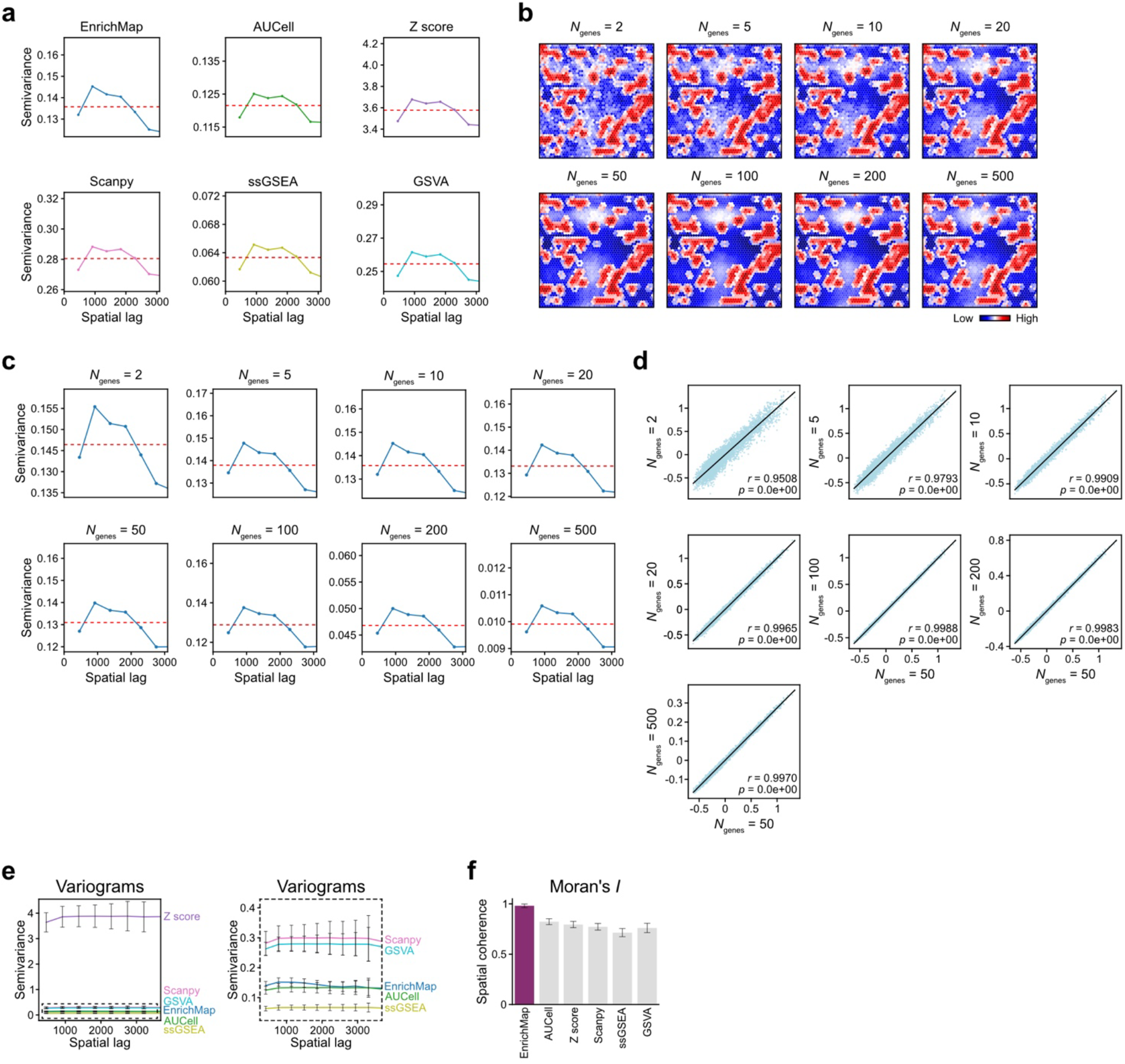
Spatial metrics of the simulated Visium-like dataset. **a**, Semivariance across spatial lags compared to the global variance across methods. **b-c**, Spatial distribution of EnrichMap scores **(b)** and their variograms **(c)** across gene sets of varying sizes. **d**, EnrichMap score correlations between the signature of 50 genes (x axis) and gene sets of varying sizes (y axis). **e**, Variograms (left) comparing different scoring methods for 100 simulations (mean ± SD). For better visibility, dashed plot (right) demonstrates variograms except for Z score (mean ± SD).

**Supplementary Fig. 2.**
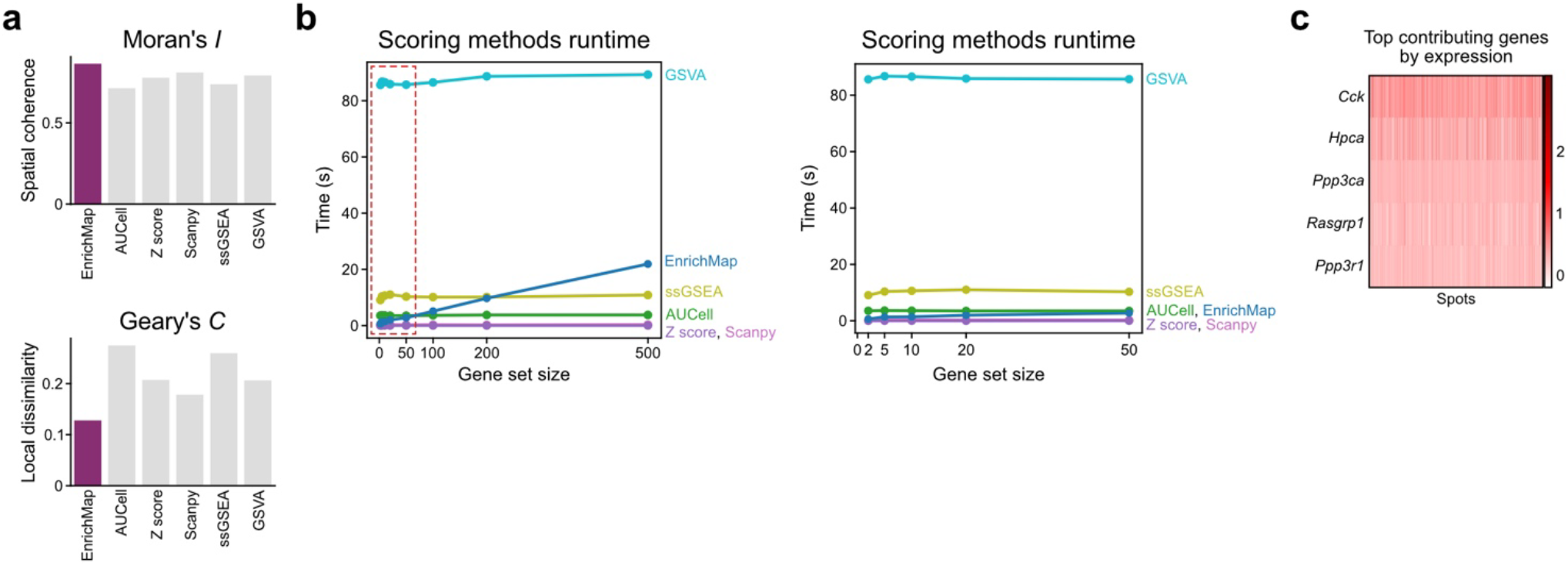
Enrichment performance in the mouse brain coronal section ST dataset. **a**, EnrichMap achieves optimal autocorrelation metrics both in Moran’s *I* (top) and Geary’s *C* (bottom). **b**, Runtime plots compare different scoring methods across varying gene set size. Left: 2-500 genes; Right: 2-50 genes for better visibility **c**, Heatmap of the top five contributing genes, sorted by average expression.

**Supplementary Fig. 3.**
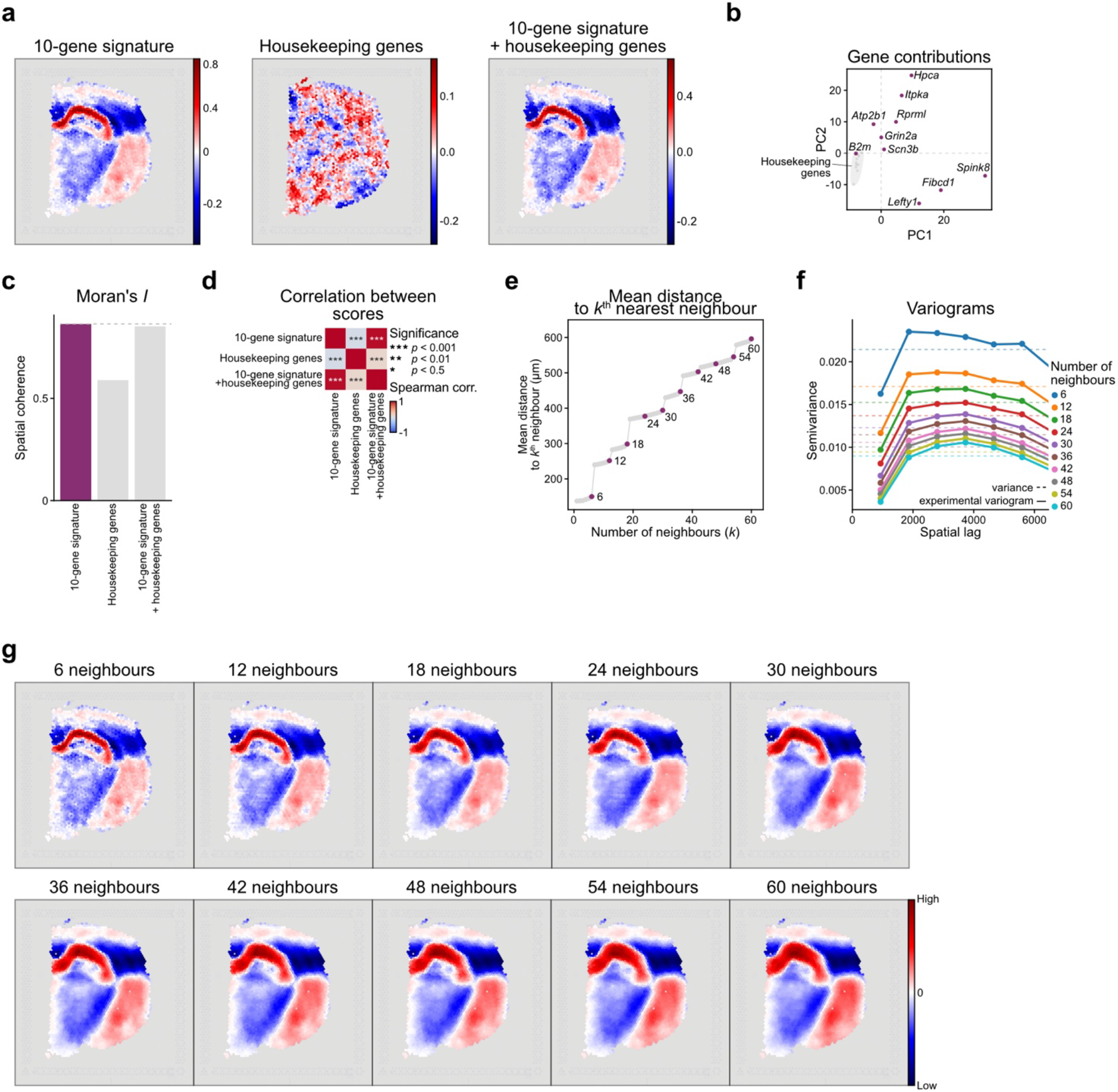
EnrichMap integrates gene-specific weights and spatial context to enhance spatial signature scoring. **a**, Spatial maps of EnrichMap scores for a 10-gene pyramidal layer signature (left), a housekeeping gene set (middle), and the combined gene list (right). **b**, Gene weights derived from the coefficient of variation suppress enrichment driven by non-informative housekeeping genes. **c**, Moran’s *I* score. **d**, Spearman correlation between 10-gene signature, housekeeping genes and combined gene set scores. **e**, Average distances (µm) to the *k*^th^ neighbour on the Visium platform, across increasing neighbour rings. **f**, Empirical variograms plotted across successive neighbour rings, capturing spatial structure in the data. **g**, Spatial EnrichMap scores computed using a range of 6^th^-order neighbourhoods.

**Supplementary Fig. 4.**
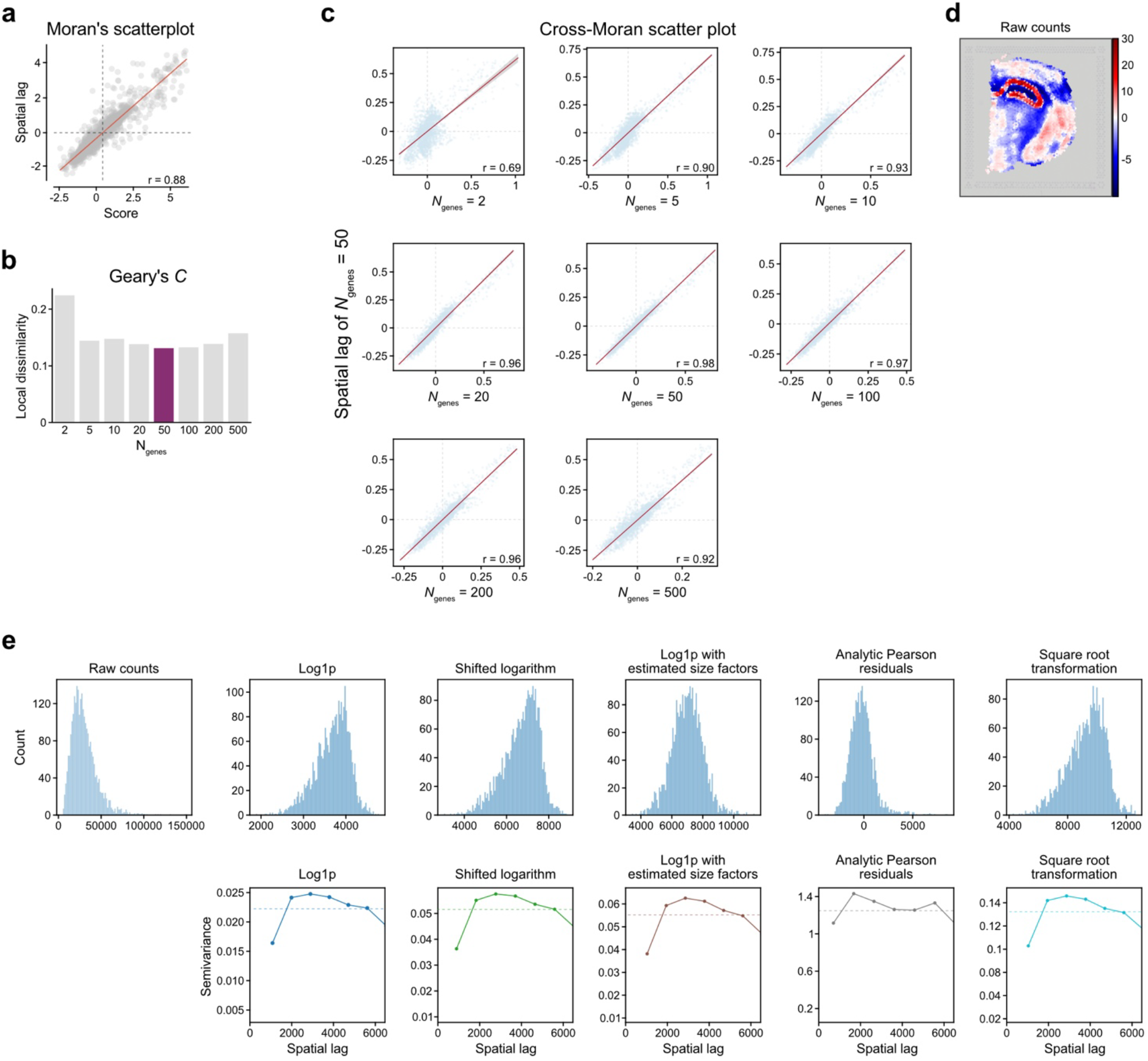
Enrichment performance in the mouse brain coronal section spatial transcriptomics dataset. **a**, Moran’s scatter plot demonstrating high correlation between the spatial lag of EnrichMap scores and raw scores. **b**, Optimal Geary’s *C* autocorrelation achieved by a signature of 50 genes. **c**, Cross-Moran plots show spatial lag of 50 genes (y-axis) and varying number of genes (2-500; x-axis), with over 93% of correlation between the optimal signature size of 50 genes and 10-100 genes. **d**, EnrichMap scores computed by raw counts. **e**, Distributions of raw and normalised counts across different methods (top) and their corresponding variograms (bottom).

**Supplementary Fig. 5.**
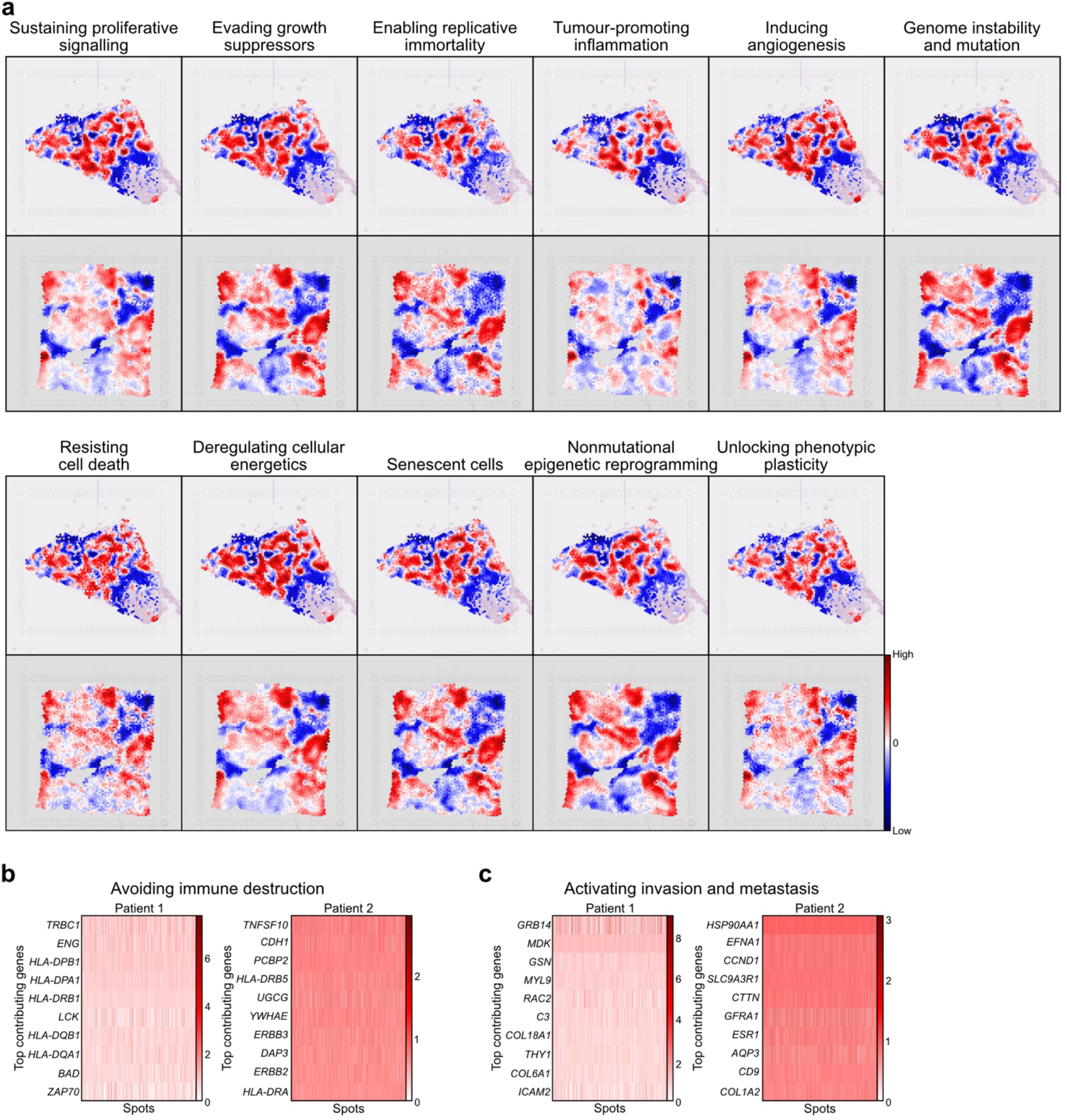
Cancer hallmarks across two breast cancer Visium slides. **a**, Cancer hallmark spatial maps across two breast cancer samples. **b-c**, Top 10 gene contributions heatmap by patient for immune evasion **(b)** and metastasis **(c)** cancer hallmarks.

